# Comparative genomics and interactions of vector thysanopterans and transmitted viruses

**DOI:** 10.1101/2025.11.21.689785

**Authors:** Michael A. Catto, Paul E. Labadie, George G. Kennedy, Alana L. Jacobson, Brendan G. Hunt, Rajagopalbabu Srinivasan

**Affiliations:** Department of Entomology, University of Georgia, Griffin, GA 30223, USA; (M.A.C.); (B.G.H); (R.S.); Department of Biological Sciences, Georgia Institute of Technology, Atlanta, GA 30332, USA; (M.A.C.); Department of Entomology and Plant Pathology, North Carolina State University, Raleigh, NC 27695, USA; (P.E.L.); (G.G.K.); Department of Entomology and Plant Pathology, Auburn University College of Agriculture, Auburn, AL 36849, USA; (A.L.J.)

**Keywords:** Core genes, insect vectors, plant viruses, protein-protein interactions, vector-virus interactions, i5K

## Abstract

We analyzed the genomes of nine thysanopteran (thrips) species, with one newly generated, and examined their relationships with 23 representative orthotospoviruses from the family *Tospoviridae* (order *Bunyavirales*). Thrips can be agricultural pests, contributing to measurable yield reductions in economically valuable crops and ornamentals. Some thrips species are confirmed orthotospovirus vectors, while most of the ∼7,000 identified species are unknown in their vector status. We conducted *in silico* protein-protein interaction predictions for several thrips proteins, including an endocuticle protein previously reported to bind to orthotospovirus glycoproteins. In most ecologically observed vector-virus pairs, the predicted protein-protein interactions were confirmed, and additional plausible vector-virus transmission interactions emerged from our analyses. These results expand our understanding of vector-virus co-evolution and highlight candidate molecular interfaces that could be targeted to disrupt virus transmission in agricultural systems.

## Introduction

Thysanopterans (thrips) are the primary vectors of orthotospoviruses, and they transmit viruses from infected to non-infected plants in a persistent and propagative manner (Ruark-Seward et al., 2020, Shrestha et al., 2019, Ullman et al., 2005). Thrips that acquire the virus exclusively during the larval stage can transmit the virus as adults (Ananthakrishnan and Annadurai, 2007, Mou et al., 2021, Wijkamp et al., 2008, Kritzman et al., 2002, Chen et al., 2006). The orthotospoviruses, in turn, have been found to exert an influence on the feeding behavior and fitness of thrips (Kindt, 2004, Stafford et al., 2011, Zhang et al., 2023b) by altering the host plant (Shalileh et al., 2016, Shrestha et al., 2012, Chen et al., 2014, Leach et al., 2019, Wu et al., 2019).

Thrips in the family *Thripidae* belong to the phytophagous suborder *Terebrantia* (order *Thysanoptera*, class *Insecta*), which is by far the most consequential group for plant-virus interactions. In contrast, thrips in the suborder *Tubulifera* cause comparatively little damage because they mainly feed on fruits and rarely attack healthy vegetative tissue. Within *Thripidae*, only members of a few genera such as *Frankliniella*, *Thrips*, and *Scirtothrips*, have been observed to transmit economically damaging viruses. Global crop losses are caused due to direct feeding and transmission of viruses by species within *Thripidae*, including the onion thrips (*Thrips tabaci*, Lindeman) (Loredo Varela and Fail, 2022), the melon thrips (*Thrips palmi*, Karny), the tobacco thrips (*Frankliniella fusca*, Hinds), the western flower thrips (*Frankliniella occidentalis*, Pergande), and the flower thrips (*Frankliniella intonsa*, Trybom) (Srinivasan et al., 2017, Adhikari et al., 2023, Wu et al., 2018, Macharia et al., 2015, He et al., 2020, Khatun et al., 2024). Some of the common susceptible and important hosts include onion (*Allium cepa* L.), garlic (*Allium sativum* L.), tobacco (*Nicotiana tabacum* L.), cotton (G*ossypium sp*. L.), chrysanthemum (*Chrysanthemum sp.* L.), and tomato (*Solanum lycopersicum* L.) (Gill et al., 2015, Loredo Varela and Fail, 2022).

More than 7,000 thysanopteran species have been described (ThripsWiki, 2005), yet only a few have annotated genomes. In this study, the newly assembled genome of the *T. tabaci* (USA) was compared with genomes of *T. tabaci* (China) (Gao et al., 2024) and with four other confirmed vector thysanopterans in *Thripidae*: *T. palmi* (Guo et al., 2020), *F. occidentalis* (Rotenberg et al., 2020), *F. fusca* (Catto et al., 2023), and *F. intonsa* (Zhang et al., 2023a). For contrast, in this study, thysanopterans of unknown vector status: the rice thrips (*Stenchaetothrips biformis*, Bagnall) (Hu et al., 2023), bean blossom thrips (*Megalurothrips usitatus*, Bagnall) (Ma et al., 2023), O*dontothrips loti* (Haliday) (Yingning et al., 2024), and the Hawaiian flower thrips (*Thrips hawaiiensis*, Morgan) (Hu et al., 2025), also were examined (**Table 1**).

**Table 1.**
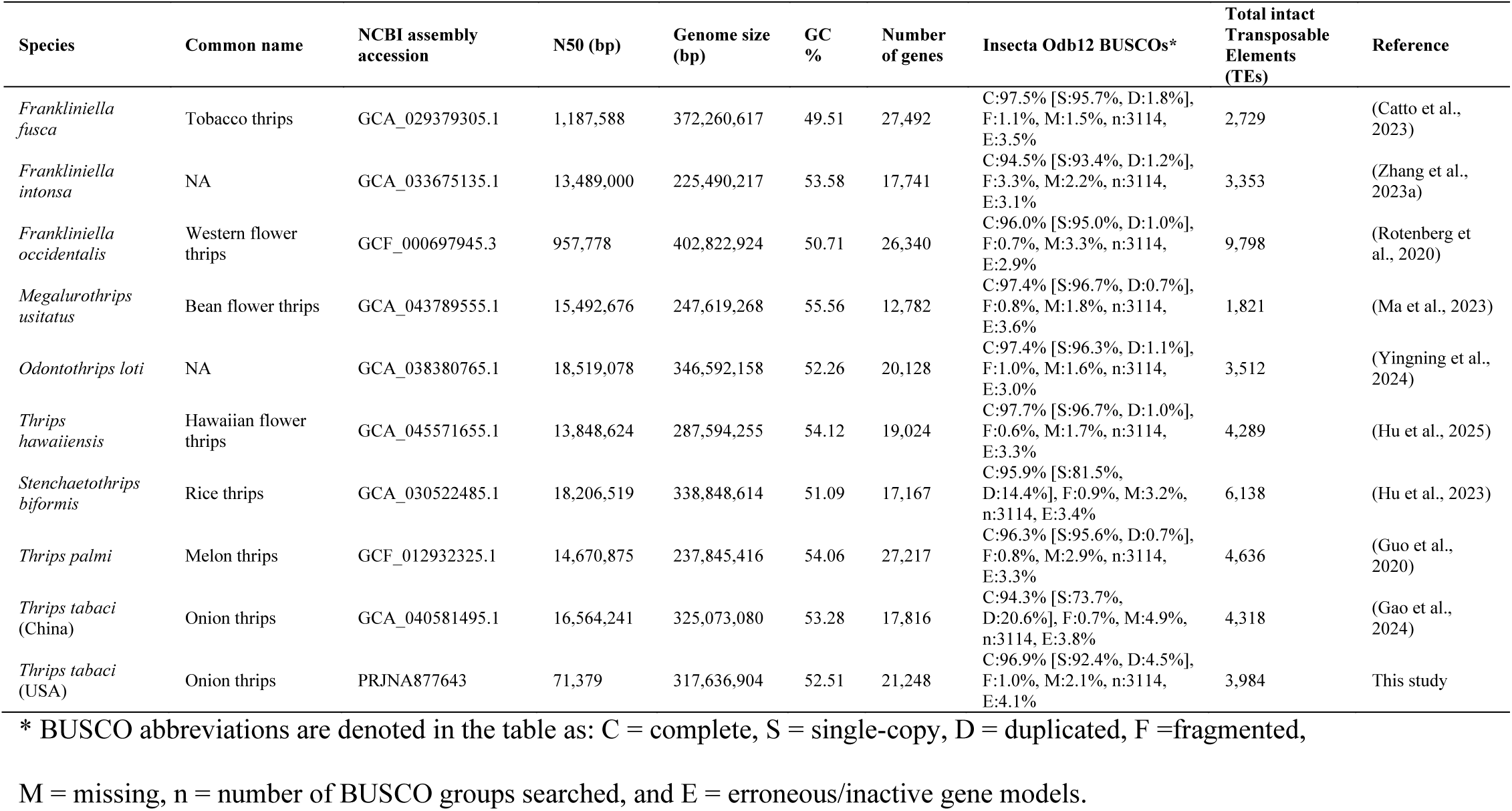
Thysanopteran genome assembly metrics.

To examine the shared evolutionary history between thysanopterans and orthotospoviruses, the genomes of 23 exemplar *Tospoviridae* species were investigated (**Table 2**). Each orthotospovirus genome comprises large (L), medium (M), and small (S) segments (Chiapello et al., 2021, Nigam and Garcia-Ruiz, 2020). The L segment encodes for the replication machinery (de Haan et al., 1991, Chao et al., 2025). The M segment encodes proteins associated with movement, pathogenicity, and vector interactions (Kormelink et al., 1992), while the S segment encodes for the nucleocapsid protein and the non-structural protein, and they are associated with virus structure and suppression of host defenses, respectively (de Haan et al., 1990). Across orthotospoviruses, five genes recur: glycoprotein (G_N_/G_C_), nucleocapsid protein (N), RNA-dependent RNA polymerase (RdRp), non-structural protein m (NSm), and non-structural protein s (NSs).

**Table 2.**
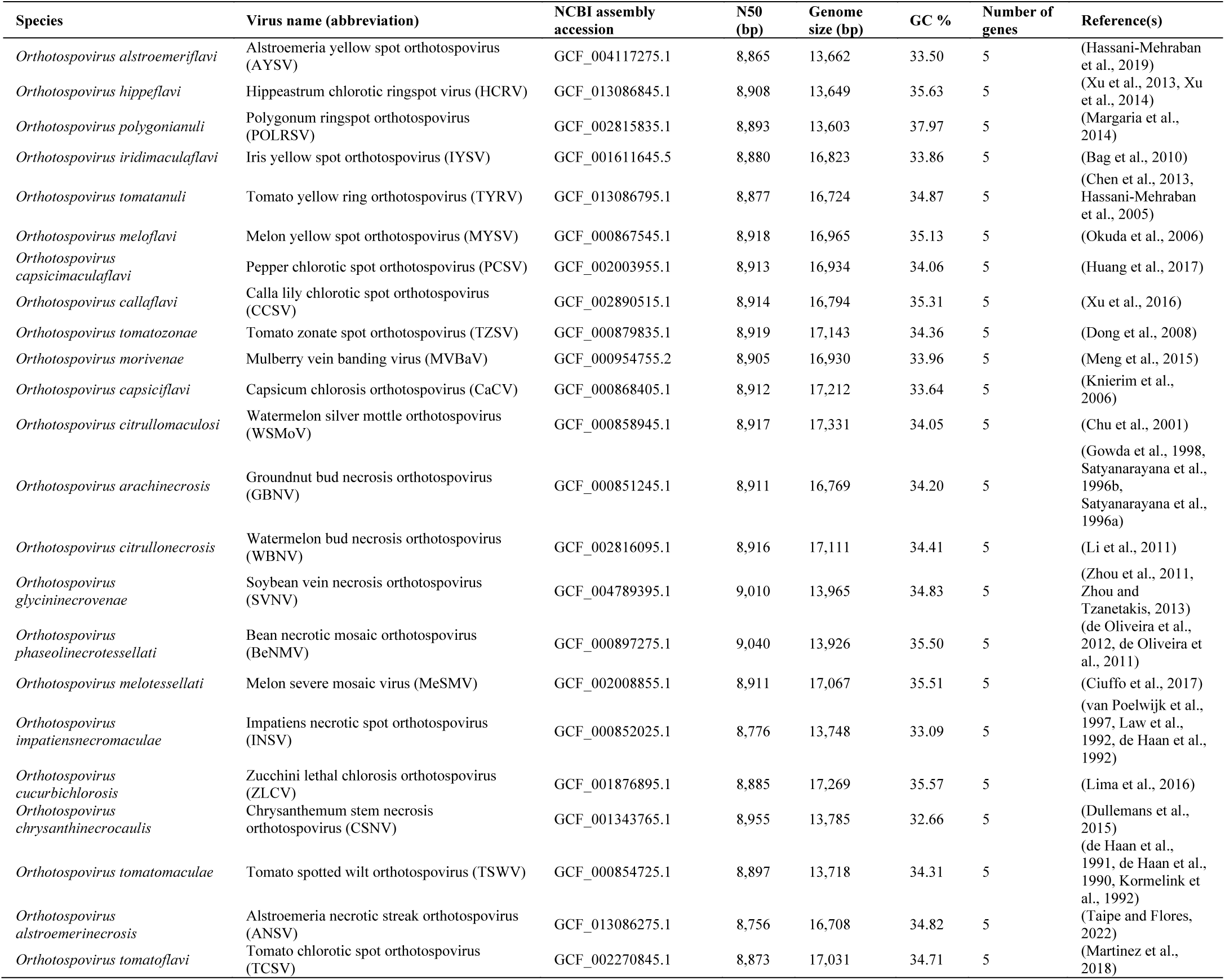
Orthotospovirus genome assembly metrics.

Our primary goal was to predict protein-protein interactions (PPIs) between a broad panel of thysanopteran vectors and orthotospoviruses. Our work started with a single experimentally confirmed thrips–virus interaction and assessed whether the same interaction context might occur in related species. By locating homologs of the five vector proteins and the virus

## Methods

### Genome acquisition and inclusion

Thysanopteran genome assemblies and annotations were downloaded from the National Center for Biotechnology Information (NCBI) RefSeq and GenBank, the Genome Warehouse (GWH), and Figshare (**Table 1**). When applicable, genome features were extracted using Another Gtf/Gff Analysis Toolkit (AGAT) v1.1.0 (Dainat et al., 2020). Orthotospovirus genome assemblies and annotations were downloaded from NCBI RefSeq (**Table 2**). The size N50 (a measure of contig length such that at least 50% of the total assembly is contained in contigs of this length or longer) and size of each assembly were determined by use of Quality Assessment Tool for Genome Assemblies (QUAST) v5.2.0 (Gurevich et al., 2013). The completeness of core genes was determined by use of Benchmarking Universal Single-Copy Orthologs (BUSCO) v5.5.0 against the Insecta Odb12 (Simao et al., 2015). Assembly reports were generated using MultiQC v1.14 (Ewels et al., 2016). Thysanopteran genome assemblies and annotations were all included for downstream analyses. Orthotospoviruses were included in downstream analyses if they had the five expected genes identified: glycoprotein (G_N_/G_C_), nucleocapsid protein (N), RNA-dependent RNA polymerase (RdRp), non-structural protein m (NSm), and non-structural protein s (NSs).

### Genome assembly, annotation, and quality control

A new *T. tabaci* genome assembly was generated for this study. 1,000 male *T. tabaci* heads from a multigeneration inbred line were used for DNA extraction to reduce heterozygosity, while providing sufficient DNA for long read sequencing (Jaron et al., 2021). PacBio library preparation was performed with the Sequel-2 chemistry kit and sequenced on two single-molecule real-time (SMRT) cells at the University of Georgia Genomics and Bioinformatics Core. Voucher specimens were retained and stored in 95% alcohol at -20C at North Carolina State University. The *T. tabaci* genome was assembled using Flye v2.9 (Freire et al., 2022) with the estimated genome size set to 400Mb, run for two iterations. Annotations from *F. fusca* (Catto et al., 2023), *F. occidentalis* (Rotenberg et al., 2020), and *T. palmi* (Chakraborty et al., 2018, Widana Gamage et al., 2018), and the *de novo* transcriptomes of *F. tritici* (Shrestha et al., 2019) and *T. tabaci* (Rosen et al., 2021) were used to train MAKER v3.01.03 (Holt and Yandell, 2011), with gene masking from Semi-HMM-based Nucleic Acid Parser (SNAP) v2013-11-29 (Korf, 2004) and AUGUSTUS v3.4.0 (Stanke and Morgenstern, 2005). The annotations were run for four iterations to refine the gene models. The genome assembly and gene models were filtered using BBMap v38.98 (Bushnell et al., 2017) to remove contamination as per the NCBI VecScreen default filtering criteria (Schaffer et al., 2018). Gene model annotations were performed using OmicBox v3.11 (Gotz et al., 2008).

*T. tabaci* forms a cryptic species complex (L1, L2, and T lineages) based upon cytochrome oxidase subunit I (COI) sequencing (Musa et al., 2023, Loredo Varela and Fail, 2022, Toda and Murai, 2007, Gawande et al., 2017, Sojnóczki et al., 2015, Li et al., 2020, Brunner and Nilsson, 2004). The mitochondrial genomes of the *T. tabaci* (USA) and the *T. tabaci* (China) were assembled using MitoHifi v3.2 (findMitoReference.py --species “*Thrips tabaci*” --min_length 14,000) based on the high quality thrips mitogenome of *Neohydatothrips gracilipes* (NCBI accession OR834984)(Ghosh et al., 2025). Minimap2 v2.28 map-pb was used to map PacBio raw reads to the reference (Li, 2018) and SAMtools v1.18 (consensus) to generate a consensus of the mapped reads (Li et al., 2009). The assemblies were aligned to *T. tabaci* COI sequences from L1 and L2 lineages (Toda and Murai, 2007) using multiple alignment using fast Fourier transform (MAFFT) v7.520 (Katoh and Standley, 2013) and a phylogenetic tree was inferred using IQ-TREE v2.3.5 with 1000 bootstrap iterations (-bb 1000 --date-options “-l 0 -u 0.001”) (Hoang et al., 2018, Minh et al., 2020, Nguyen et al., 2015). Thysanopteran transposable elements (TEs) were annotated by using the Extensive *de novo* TE Annotator (EDTA) tool v2.2.2 (Ou et al., 2019) and associated panEDTA tool (Ou et al., 2024). The EggNOG-mapper v2.1.9 (Huerta-Cepas et al., 2017, Huerta-Cepas et al., 2019) and DIAMOND v2.1.9 (Buchfink et al., 2021) were used to assign functional annotations to each of the species gene models (emapper.py --override --cpu 0 -m diamond --decorate_gff yes).

### Orthogroups detection and phylogenetic tree construction

Orthologs were determined by using Orthofinder v2.5.5 (Emms and Kelly, 2015, Emms and Kelly, 2019) with MAFFT v7.520 (Katoh and Standley, 2013) and FastTree v2.1.11 (Price et al., 2010). Outgroup proteomes, being the pea aphid (*Acyrthosiphon pisum*, Harris) (International Aphid Genomics, 2010, Li et al., 2019) and the sweetpotato whitefly (*Bemisia tabaci*, Gennadius) (Chen et al., 2016, ROTHAMSTED, 2022, Thao and Baumann, 2004, Thao et al., 2004), were used for tree building, but were not included in downstream analyses. Alignment statistics were calculated using the AliStat tool (Wong et al., 2020). The IQ-TREE v2.3.5 (Hoang et al., 2018, Minh et al., 2020, Nguyen et al., 2015) was used to construct a species tree from 1000 bootstrap iterations (-bb 1000 --date-options “-l 0 -u 0.001”) from the OrthoFinder2 species tree alignment fasta files. Figtree v1.4 (https://github.com/rambaut/figtree/) was used for visualization of the resulting tree file.

Roary v3.13, using the following parameters (-ap -i 25 -e -mafft), was used to find the representative sequences of the super pan-genome (Page et al., 2015). PanTools v3.4, using the following parameter (group --intersection-rate=0.001 --similarity-threshold=8 --mcl-inflation=1 - -contrast=0), was used to determine the Heaps law α (Jonkheer et al., 2022, Sheikhizadeh et al., 2016, Jonkheer et al., 2025, Sheikhizadeh Anari et al., 2018). Pan-genomes are considered open (Heaps law α ≤ 1) or closed (Heaps law α > 1), which is used to determine if the discovery of new gene clusters is expansive or constant, respectively (Tettelin et al., 2005).

### Molecular evolution

Orthologr (Drost et al., 2015, Buchfink et al., 2021, Needleman and Wunsch, 1970, Suyama et al., 2006, Zhang et al., 2006, Comeron, 1995) was used to to determine initial pairwise comparisons of positive selection of other thrips species relative to the USA and Chinese lineages of *T. tabaci*. To compare the strength of natural selection between confirmed virus-transmitting vectors and unknown vectors across the broader thysanopteran dataset, we extracted single-copy orthogroups with OrthoFinder2 (Emms and Kelly, 2015, Emms and Kelly, 2019) and generated codon alignments using PAL2NAL v14.1 (Suyama et al., 2006). The resulting alignments, together with the species tree inferred by OrthoFinder2, served as input for HyPhy v2.5.33 Adaptive Branch-Site Random Effects Likelihood (aBSREL) (Smith et al., 2015). aBSREL tested each branch for episodic diversifying selection relative to the *T. tabaci* USA or Chinese lineages, the non-reference *T. tabaci* lineage was excluded from the tests. aBSREL was executed with its default parameters to test all branches, and branch-wise p-values were Holm-Bonferroni corrected (α = 0.05). Branches were labeled based on confirmed vector and unknown vector status based on ThripsNET. (n.d.). Key server – Thrips of North America. Retrieved August 9, 2025, from https://thripsnet.zoologie.uni-halle.de/key-server-neu/data/0a08090e-0e03-4a0e-8502-070105080e05/media/Html/index.htm. HyPhy v2.5.33 RELAX (Wertheim et al., 2015) was applied to the same alignments and tree to assess whether positive selection pressure on particular lineages were intensified (k > 1) or relaxed (k < 1) compared with the background, again using Holm-Bonferroni correction.

### Predicting protein-protein interactions

Sequences of interest XP_017786818.1 (cell surface glycoprotein 1), XP_018334183.1 (endocuticle structural glycoprotein SgAbd-2-like), XP_019753975.1 (peptidyl-prolyl cis-trans isomerase), XP_019767728.1 (enolase), XP_023718907.1 (ATP synthase subunit alpha, mitochondrial), and XP_022906571.1 (endocuticle structural glycoprotein SgAbd-2-like) (Badillo-Vargas et al., 2019, Maurastoni et al., 2023), were downloaded from NCBI. These sequences were then run through the Protein Basic Local Alignment Search Tool (BLASTP) (Camacho et al., 2009) to find sequences with similarity in the thysanopteran proteomes. Using the list of hits from BLASTP, OrthoFinder2 orthogroups were selected for downstream protein-protein interaction testing, which yielded orthogroups: endocuticle structural glycoprotein (OG0007628), spliceosome-associated protein (OG0008117), V-type proton ATPase (OG0008221), peptidyl-prolyl cis-trans isomerase (OG0009306), and RING-type E3 ubiquitin-protein ligase (OG0009466). Orthotospovirus orthogroup OG0000004 was selected, as it contained the structural glycoproteins (G_N_/G_C_) of all 23 orthotospoviruses. SpeedPPI v20230913 “create_ppi_some_vs_some.sh” was used to predict protein-protein interactions using default cutoff scores of pdockq = 0.5 (Bryant et al., 2022). Proteins were visualized using the ChimeraX v1.6.1 (Goddard et al., 2018, Pettersen et al., 2021, Meng et al., 2023). To compare predicted protein-protein interactions, we used Root Mean Square Deviations (RMSDs) calculated with TM-align v20180426 (Zhang and Skolnick, 2005), which quantifies the average atomic distance between aligned 3D structures.

## Results

### Gene orthology and phylogenetics of thysanopterans and orthotospoviruses

Comparative genomic analyses of nine thysanopteran species revealed evolutionary relationships that provide context for interpreting patterns of vector competence and genome composition. A species tree constructed from 1,067 single-copy orthogroups, resolved well-supported relationships among these species (**Fig. 1A; Additional File 2: Tables S1 – S4**). This phylogenetic framework facilitated comparisons of other genomic features, such as transposable element (TE) composition, which showed lineage-specific expansions in several clades (**Additional File 1: Fig. S1; Additional File 2: Table S5**). Across the reconstructed phylogeny, both putative vector and non-vector taxa are interspersed rather than forming separate clades. This pattern indicates that traits associated with vector competence have likely emerged, or been lost, independently in multiple lineages. Many of the thysanopterans examined have not been experimentally confirmed as virus transmitters, but could be problematic in introduced agricultural settings due to the uncertainty surrounding their role in virus transmission.

**Figure 1.**
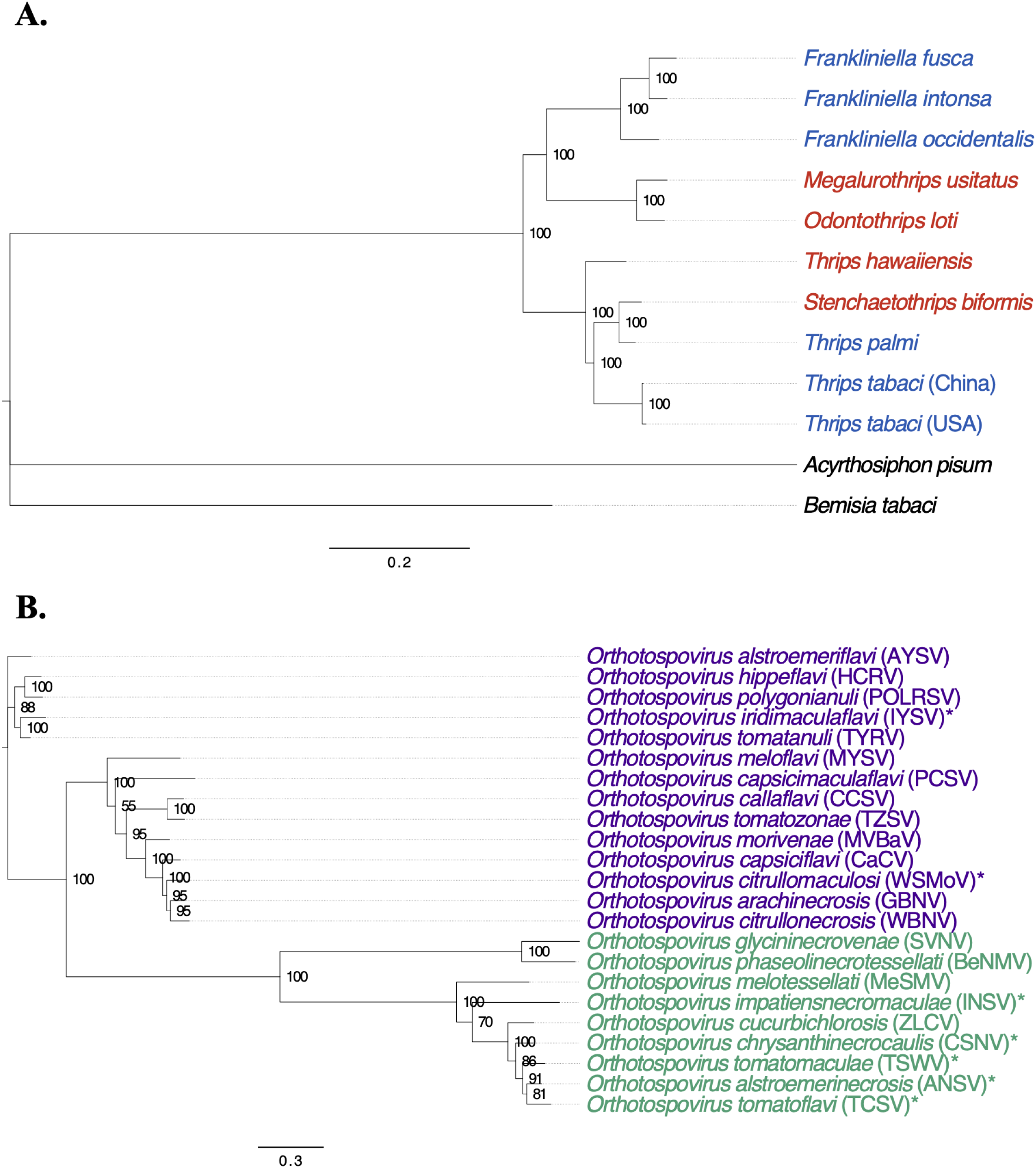
Phylogenetic trees of thysanopterans and orthotospoviruses. (**A.**) Species tree of thysanopterans constructed from 373 single-copy orthogroups with pea aphid (*A. pisum*) and sweetpotato whitefly (*B. tabaci*) included as outgroup taxa. Bootstrap support values are based on 1,000 ultrafast replicates in IQ-TREE. While the thysanopteran dataset contained 1,067 single-copy orthogroups, the tree shown here reflects the inclusion of outgroups. Red and blue indicate unknown and confirmed vector status based on ThripsNET, respectively. (**B.**) Orthotospovirus species tree was constructed from five single copy orthogroups using OrthoFinder2. Bootstrap values come from 1,000 iterations of IQ-TREE. American and Eurasian clades are indicated by green and purple respectively, and serotypes denoted with an asterisk (Cheng et al., 2021, Feng et al., 2023, Chao et al., 2025).

A complementary phylogenetic tree of orthotospoviruses was constructed using five single-copy orthogroups across 23 species (**Fig. 1B; Additional File 3: Tables S6 – S10**). The resulting topology revealed strong support for major clades (American and Eurasian) corresponding to established serogroups (Cheng et al., 2021, Feng et al., 2023, Chao et al., 2025). The relatively conserved nature of these virus proteins across divergent lineages supports their functional importance and cross-species relevance in vector interactions.

### Molecular evolution

To assess patterns of adaptive change across the order *Thysanoptera*, we reconstructed a species-level phylogeny using single-copy orthologs and then applied branch-site codon models to each orthogroup (**Additional File 4: Table S11**). Although the positive selection analysis encompassed representatives from all major thrip genera, the two geographically distinct lineages of *Thrips tabaci* (USA and China) were examined as independent focal branches because *T. tabaci* is a well-studiedphytophagous member of the suborder *Terebrantia* and serves as a proxy for many vector species within the family (**Additional File 4: Table S12**). Genes identified by the positive-selection tests were compared, revealing both overlapping and lineage-specific subsets that show elevated dN/dS ratios (> 1) –a pattern indicative of episodic positive selection in each lineage (**Fig. 2A**; **Additional File 1: Fig. S2; Additional File 4: Table S13**). We compared the selection intensity on foreground branches representing confirmed virus-transmitting thrips versus background branches comprising unknown vector taxa. With respect to *T. tabaci* (USA), 98/973 orthogroups showed a statistically significant intensification (k > 1, p ≤ 0.05) and 165/973 displayed relaxation (k < 1, p ≤ 0.05) (**Additional File 4: Table S14**).

**Figure 2.**
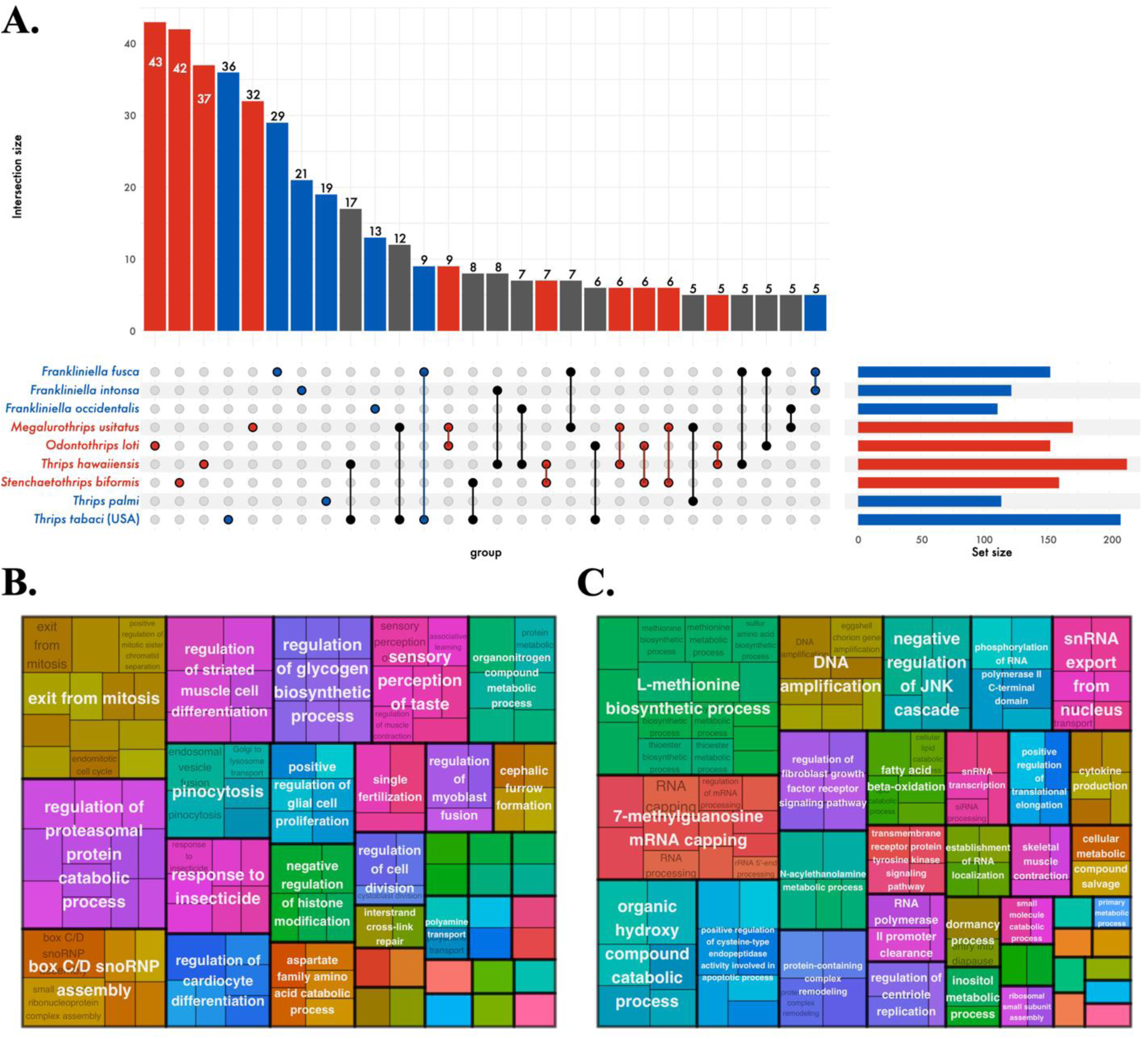
Overlap and Gene Ontology (GO) enrichment of orthogroups under positive selection across thysanopterans. (**A.**) UpSet plots illustrate the number and intersection of orthogroups showing signatures of positive selection (dN/dS > 1, based on aBSREL analysis) across thysanopteran species. Minimum size was set to 5 for visualization purposes, the full list of overlapping genes can be found in **Additional File 4: Table S13**. Vertical bars indicate the number of orthogroups shared across species or unique to one lineage, while horizontal bars show the total number of positively selected orthogroups per species. Red and blue indicate unknown and confirmed vector status based on ThripsNET, respectively. GO enrichment of (**B.**) confirmed and (**C.**) unknown thysanopteran vectors of orthotospoviruses. Results based on selection testing that excludes *T. tabaci* (China) lineage.

Gene Ontology (GO) enrichment analyses were conducted (**Additional File 1: Fig. S3; Additional Files 5 & 6: Tables S15 & S16**). Genes subject to positive selection in one or more confirmed vector species were enriched for processes associated with cell cycle regulation (e.g., exit from mitosis, regulation of mitotic sister chromatid segregation), protein and RNA complex assembly (e.g., box C/D snoRNP assembly, regulation of glucan biosynthetic process), proteasomal protein catabolic processes, and more specialized cellular activities including response to insecticide, sensory perception, regulation of muscle cell differentiation, polyamine transport, and developmental patterning (e.g., imaginal disc-derived wing hair orientation, germ-band extension; **Fig. 2B; Additional File 5: Table S15**). In contrast, unknown vectors showed enrichment for processes linked to RNA metabolism (e.g., RNA capping, mRNA processing, snRNA export), DNA replication and amplification, lipid and small-molecule metabolic processes (e.g., fatty acid beta-oxidation, inositol metabolism), immune-related regulation (e.g., negative regulation of inflammatory response, cytokine production), and diverse sensory and developmental pathways (e.g., dormancy process, detection of temperature stimulus; **Fig 2C; Additional File 5: Table S15**).

### Vector status related to predicted protein-protein interactions

Protein-protein interactions (PPIs) between insect vectors and plant viruses play a critical role in virus acquisition, retention, and inoculation. In orthotospoviruses such as the TSWV glycoprotein (G_N_/G_C_) mediates attachment and entry into host insect tissues, particularly within the midgut and salivary glands. Prior molecular studies in *F. occidentalis* have identified direct interactions between TSWV G_N_/G_C_ and cuticular proteins of the vector, suggesting a mechanistic basis for tissue tropism and vector specificity (Badillo-Vargas et al., 2019, Maurastoni et al., 2023).

To explore whether such interactions show signatures associated with vector status, *F. occidentalis* proteins experimentally confirmed to bind TSWV G_N_/G_C_ (Badillo-Vargas et al., 2019, Maurastoni et al., 2023) were used to identify single copy orthogroups of interest across a representative set of thysanopteran species (**Additional File 7: Table S17**). Predictive modelling was then applied to assess PPIs between these insect homologs and orthotospovirus G_N_/G_C_ homologs, using the experimentally validated interactions as templates. Examples of predicted interactions between thysanopteran V-type proton ATPases with INSV G_N_/G_C_ and TSWV G_N_/G_C_ are shown in **Fig. 3** and **Additional File 1: Fig. S3**, respectively. Across the five models shown in **Fig. 3**, the number of contacts ranged from 919 to 1,327 (mean ≈ 1,172 ± 148), with consistently high confidence scores (average pLDDT ≈ 70.8 ± 2.1 and pDockQ ≈ 0.718 ± 0.004) (**Additional File 7: Table S18**). Detailed PPI hydrogen contact points are provided in **Additional File 7: Table S19**.

**Figure 3.**
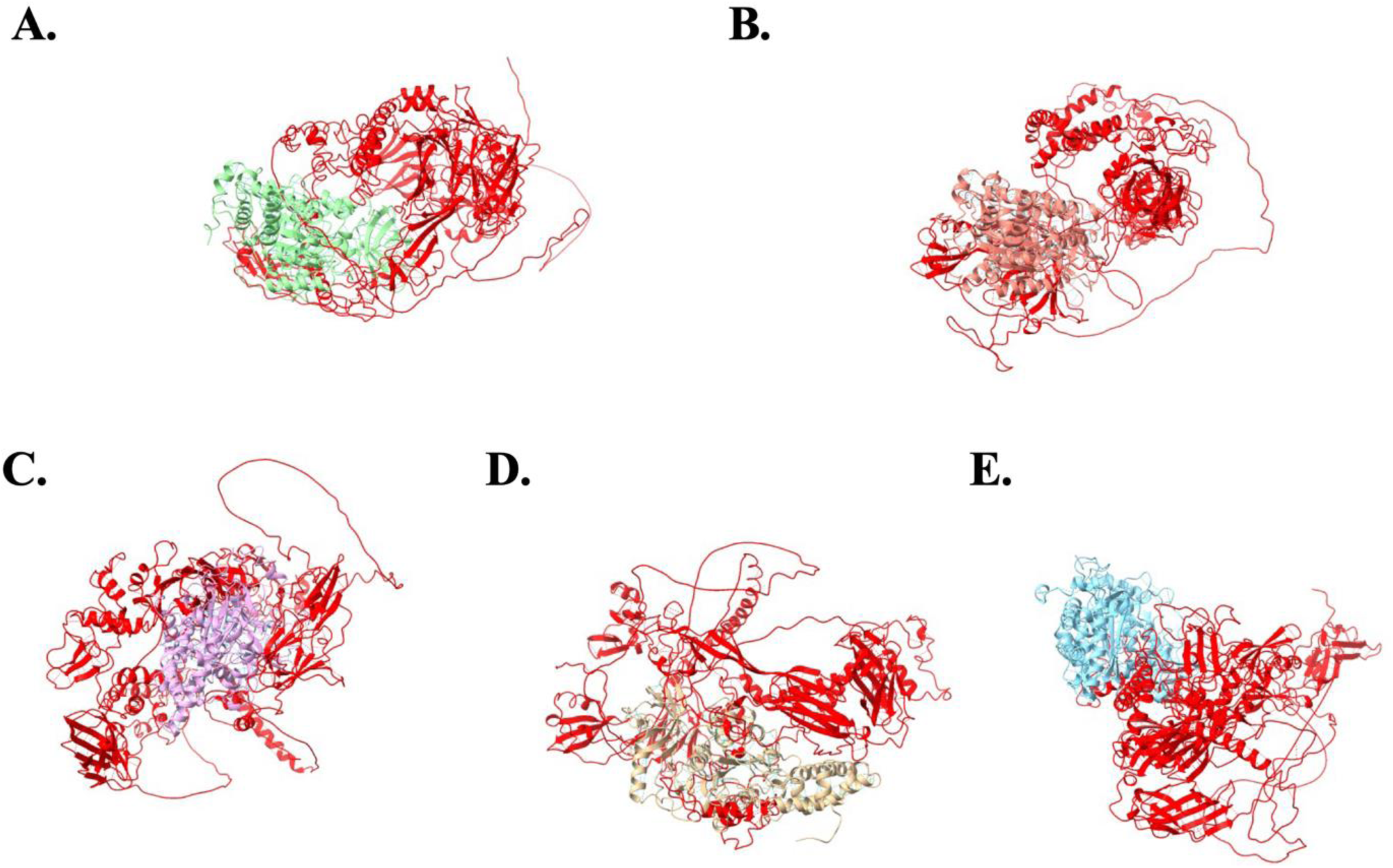
Predicted protein-protein interactions (pDockQ ≥ 0.7) between thysanopteran proteins (green, coral, magenta, tan, cyan) and orthotospovirus proteins (red), based on hydrogen bonding (blue). All models were aligned to the *T. tabaci* (USA) – *O. impatiensnecromaculae* glycoprotein subchain. Protein-protein interaction examples from (**A.**) *T. tabaci* (USA), (**B.**) *T. tabaci* (China), (**C.**) *S. biformis*, (**D.**) *T. hawaiiensis*, and (**E.**) *M. usitatus*.

The thysanopteran single copy orthogroups are: endocuticle structural glycoprotein (OG0007628), spliceosome-associated protein (OG0008117), V-type proton ATPase (OG0008221), peptidyl-prolyl cis-trans isomerase (OG0009306), and RING-type E3 ubiquitin-protein ligase (OG0009466). Orthotospovirus single copy orthogroup is the structural glycoprotein (G_N_/G_C_) (OG0000004). A comparison of all tested PPIs in relation to confirmed and unknown vector status is presented in **Fig. 4A** and **Additional File 7: Table S20**. Overall, positive PPIs were observed in 59% of species of unknown vector status interactions (595/1005) and 58% of confirmed vector interactions (84/145). Statistical tests indicated no significant difference between these groups (χ² = 0.04, df = 1, p = 0.84; Fisher’s exact p = 0.79), suggesting that predicted interaction frequency was broadly similar regardless of vector status.

**Figure 4.**
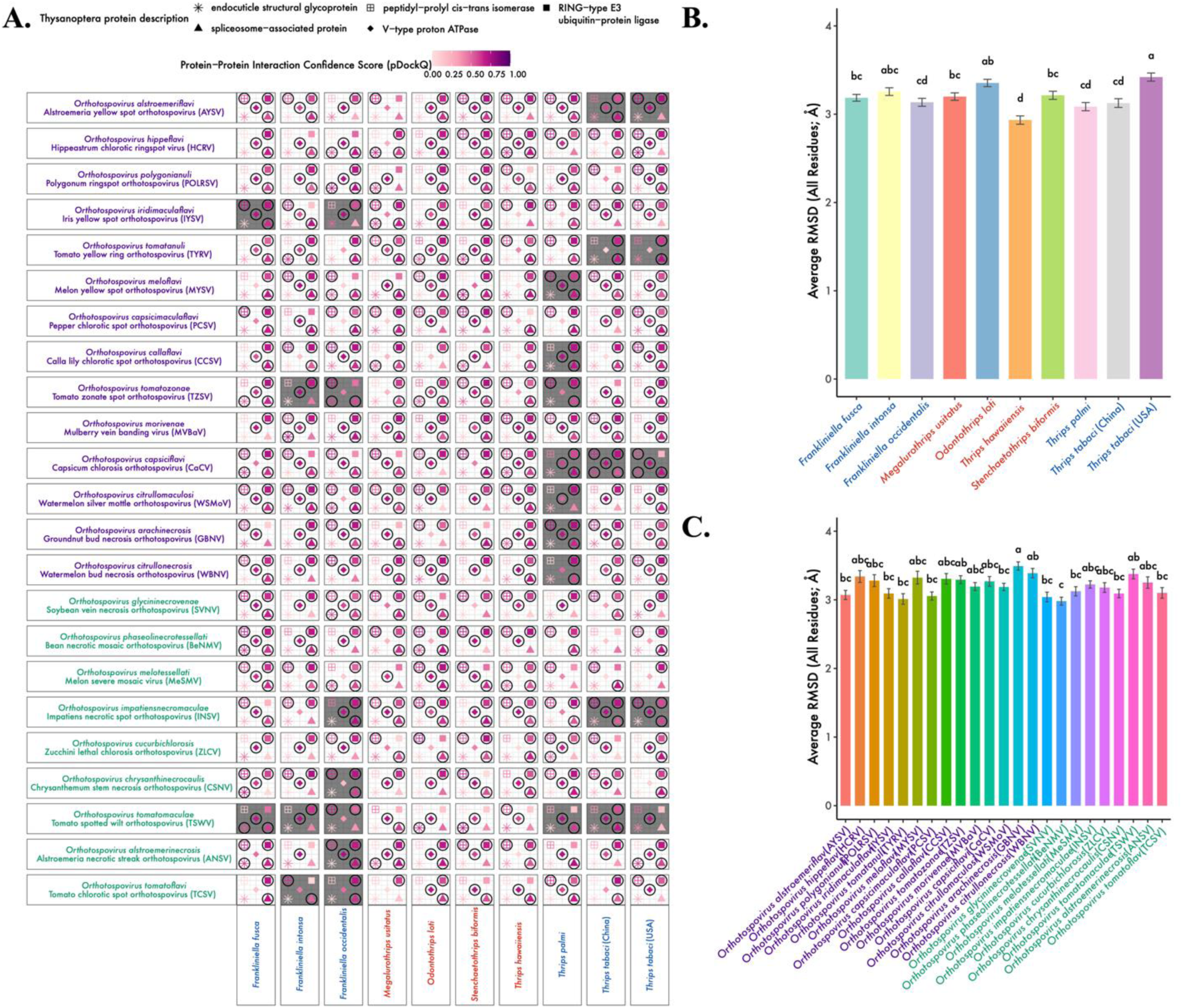
Vector status and protein-protein interaction comparisons and model alignment. (**A.**) Protein-protein interaction predictions were inferred via SpeedPPI/AlphaFold2 modeling. Each shape represents the presence of a predicted protein-protein interaction between a thysanopteran protein and orthotospovirus protein from one of five orthogroup pairs. Shapes with a circle around them passed a model confidence threshold of pDockQ ≥ 0.5. Tiles show vector status as confirmed (grey) and unknown (white) for each orthotospovirus. (**B.**) Thysanopteran and (**C.**) orthotospovirus average root-mean-square deviation (RMSD; Å) from TM-align structural superpositions of all residues across high-confidence matches (coverage ≥ 0.4, alignment length ≥ 100 residues). Lower RMSD values indicate greater structural similarity. Red and blue indicate unknown and confirmed vector status based on ThripsNET, respectively. Orthotospovirus American and Eurasian clades are indicated by green and purple, respectively (Cheng et al., 2021, Feng et al., 2023, Chao et al., 2025).

Pairwise average root-mean-square deviation (RMSD; Å) comparisons between atomic structures of PPI were compared to test conservation of protein folding. Across all comparisons, RMSD values were generally low (median ≈ 4.24 Å; average ≈ 4.4 Å), with nearly 39% of alignments below 3 Å and ∼23% below 2 Å (**Additional File 7: Table S21**). Comparisons between the folding patterns within species found conservation of models with the average RMSD being ∼3.2 Å for both thysanopterans and orthotospoviruses (**Figs. 4B & 4C; Additional File 7: Table S22**).

## Discussion

A comparative-genomic profile of nine thysanopteran species was generated and examined how their proteins might interact with orthotospoviruses in this study. Computational testing of identified protein-protein interactions (PPI) between *T. tabaci* and several orthotospoviruses, including AYSV, CaCV, INSV, IYSV, TYRV, and TSWV, was consistent with previous reports of observed vector-virus pairs (Cabrera-La Rosa and Kennedy, 2007, Ghotbi and Baniameri, 2006, Diaz-Montano et al., 2010, Birithia et al., 2013, Oliver and Whitfield, 2016, Mortazavi and Aleosfoor, 2015, Riley et al., 2011, Widana Gamage et al., 2018, Loredo Varela and Fail, 2022). Likewise, the other confirmed vectors each had PPI with orthotospoviruses they have been observed to transmit (Chiaki et al., 2020, Mou et al., 2021, Ghosh et al., 2021, Oliver and Whitfield, 2016, Adachi-Fukunaga et al., 2020, Chen et al., 2020, Riley et al., 2011, LaTora et al., 2022, Srinivasan et al., 2012, Shrestha et al., 2017, Lai et al., 2021, Stumpf and Kennedy, 2007, Stumpf and Kennedy, 2004, Li et al., 2023a, Khatun et al., 2024, Wijkamp, 1995, Nagata et al., 2004, Nagata and de Ávila, 2000, Birithia et al., 2013, He et al., 2020). Similiarly, we found that the unknown vector species that were tested in this study may have the potential to interact with orthotospoviruses, even though they have not been observed to do so in nature (Moritz et al., 2009).

This investigation focused on a set of orthogroups previously linked to orthotospovirus entry, specifically endocuticle glycoproteins, spliceosomes, ATPases, peptidyl-prolyl isomerases, and ubiquitin-related proteins, that interact with the glycoprotein in the *F. occidentalis*–TSWV system (Badillo-Vargas et al., 2019, Maurastoni et al., 2023, Tomitaka, 2019). PPI predictions yielded numerous high-confidence contacts across these examined vector-virus pairs. While the predicted PPIs may reflect species-specific differences, further experiemental testing could provide more insight into these interactions. The differences between PPI observations and observed interaction may stem from several, not mutually exclusive, scenarios such as (i) spatial isolation between plant, vector, and virus, (ii) molecular barriers that prevent binding, (iii) ancesteral potentials that were lost as lineages diverged, or (iv) genuine but undocumented interactions. Taken together, the coexistence of conserved entry factors and lineage-specific divergences suggests that orthotospovirus transmission is shaped both by deep evolutionary constraints and by more recent adaptive shifts within thrips lineages.

Intracellular chaperones such as glycoproteins emerged as frequent predicted partners of virus structural glycoprotein (G_N_/G_C_), indicating a potential role in post-entry virus processing or replication (Khan et al., 2023). These proteins are frequently utilized by other viruses and have been identified in tissues relevant to virus propagation, such as the midgut and salivary glands (Khan et al., 2023). Moreover, positive selection was detected in a subset of immune-related orthologs, implying a co-evolutionary system with persistent propagative viruses. Notably, some unknown vector species lack compatible integrin or endocuticle variants (e.g., *Frankliniella tritici*; (Shrestha et al., 2019)), reinforcing the idea that subtle protein-sequence differences can influence vector competence.

Orthotospoviruses can be acquired by thrips and after acquisition, G_N_ and G_C_ glycoproteins facilitate attachment to and entry into the insect’s midgut cells, ensuring successful transmission (Sin et al., 2005). Phylogenetic analyses resolve orthotospoviruses into seven canonical serogroups, with a few divergent isolates occasionally placed in additional provisional groups (Zhang et al., 2021). Because serogroup membership does not strictly limit vector competence, the two most ubiquitous vectors, *F. occidentalis* and *T. tabaci*, have been shown to transmit at least twelve orthotospovirus species across three serogroups, including TSWV, GRSV, and IYSV (Nagata et al., 2004).

Since the virus entry machinery imposes few intrinsic constraints on vector compatibility, extrinsic ecological variables: host-plant preference, geographic range, and seasonal abundance, become the primary determinants of transmission competence. The introduction of TSWV into the California Central Valley, where *F. occidentalis* and *T. tabaci* were already abundant, triggered severe crop losses from 2005-2006 that cost processing tomato growers millions of dollars, largely because dense thrips populations and plant reservoirs allowed TSWV to spread rapidly (Batuman et al., 2020). A similar situation could potentially occur in other vector-virus systems based on PPI predictions. Whenever a newly introduced orthotospovirus meets an existing thrips community, proactive monitoring of vector populations and systematic sampling of ornamental and crop reservoirs become essential for early detection and containment (Chitturi et al., 2018). Likewise, when thrips vectors are introduced into new ecosystems, baseline surveys of thrips species and regular checks of host-plant availability are crucial to prevent unexpected virus outbreaks.

Expanding these approaches to additional species and testing the function of candidate proteins will help clarify the molecular basis of transmission and could inform strategies to reduce the spread of orthotospoviruses in crops. Expanding genomic sampling to additional populations of *T. tabaci* (Jouraku et al., 2025), *F. occidentalis* (Song et al., 2024), or understudied taxa such as *Dendrothrips minowai* (Xiu et al., 2025) would further our understanding of thysanopteran diversity. A pangeomic resource with more thrips species could also help to further shed light on these vector-virus interactions (He et al., 2025, Jonkheer et al., 2025, Matthews et al., 2024, Leonard et al., 2023, Li et al., 2023b, Cochetel et al., 2023, Lobb et al., 2023, Tetz and Tetz, 2020, Khan et al., 2020, Golicz et al., 2020, Aherfi et al., 2018, Muhammad et al., 2023, Ding et al., 2018, Sheikhizadeh Anari et al., 2018).

## Supplementary files

Please see additional files for Tables S1 – S22 and Figures S1 – S4.

## Data availability

The *Thrips tabaci* genome assembly and associated gene models are publicly available through NCBI under BioProject accession PRJNA877643, with the assembly accession JAUNZC000000000. The corresponding raw sequencing reads have been deposited under BioSample accession SAMN30716846 and SRA accession SRR21469203. The *Thrips tabaci* mitochondrial genomes, thysanoptera transposable element annotations, the orthotospovirus pangenome, the thysanoptera and orthotospovirus orthogroups, and predicted protein-protein interactions and associated submission scripts have been deposited to the FigShare repository in following link https://figshare.com/s/f9158870098b39057b68.

## Acknowledgements

We would like to thank the University of Maryland Brain & Behavior Institute – Advanced Genomic Technologies Core for conducting the sequencing of our samples.

## Author contributions

A.L.J. and R.S. obtained funding; P.E.L, G.G.K., and A.L.J. reared insect cultures; P.E.L. performed extractions; M.A.C. executed data analysis pipelines; M.A.C. worked on data management; B.G.H provided oversight for computational and statistical analyses; M.A.C, B.H.G., and R.S wrote the manuscript; All authors contributed to further edits and revisions.

